# DMPKformer: An Interpretable Multimodal Deep Learning Framework for Reliable ADMET Property Prediction

**DOI:** 10.64898/2026.05.28.728612

**Authors:** A.S. Ben Geoffrey, Abhishek Singh, Sowmya Kanchan, Samir Anapat, Kishan Gurram, Nagaraj M Kulkarni

## Abstract

Accurate prediction of absorption, distribution, metabolism, excretion, and toxicity (ADMET) properties remains a critical challenge in drug discovery. Traditional single modality approaches often fail to capture the complex, multi-scale relationships governing molecular behavior across physicochemical, structural, and pharmacokinetic dimensions. In this work, we propose a multi-modal deep learning framework that integrates complementary molecular representations, MACCS fingerprints, molecular graphs, and physicochemical descriptors to achieve robust ADMET property prediction. Each modality is modeled using a specialized neural subnetwork tailored to its structural characteristics: a self-attention–based Transformer encoder for MACCS fingerprints, a Graph Attention Network (GAT) for molecular graph representations, and a tanh-activated multilayer perceptron for RDKit-, PaDEL-, and Mordred-derived descriptors. Each modality is independently trained for binary classification, and latent embeddings extracted from internal layers serve as transferable molecular representations. These embeddings are subsequently fused and fine-tuned via a tanh-activated dense network and shared prediction head to form a unified ADMET predictor. The proposed framework achieves competitive performance across multiple TDC ADMET benchmarks while providing enhanced interpretability through modality-specific attention mechanisms. In addition, the incorporation of latent-space out-of-distribution (OOD) confidence estimation enables identification of high-confidence operating regions, improving the reliability and practical applicability of the framework for molecular property prediction in drug discovery workflows.

## 1. Introduction

ADMET property prediction is an indispensable component of modern drug discovery pipelines, enabling early identification of liabilities related to pharmacokinetics and toxicity before experimental validation. Despite substantial progress in computational chemistry, predictive modeling of ADMET endpoints remains challenging due to the inherent heterogeneity and complexity of molecular data. Molecules exhibit multi-scale structure–property relationships, ranging from atom-level connectivity and local substructures to global physicochemical characteristics, that are not fully captured by any single data modality.

Recent advances in deep learning for molecular representation have introduced architectures tailored to specific input formats. Graph neural networks (GNNs) efficiently encode topological and relational information from molecular graphs, while Transformer-based models capture global dependencies across fingerprint features, including large-scale molecular language models such as MolFormer [1-6]. Similarly, descriptor-based neural networks can leverage curated physicochemical features such as hydrophobicity, polar surface area, and molecular weight [7-8]. However, single-modality models are inherently limited in their representational capacity, as they often overlook cross-modal correlations and complementary information across feature spaces [9-12].

To address this limitation, we propose a multimodal ADMET prediction framework that integrates three complementary molecular representations: (1) MACCS structural fingerprints, which encode fragment-level structural patterns; (2) graph-based molecular representations, which capture atomic connectivity and local neighborhood interactions; and (3) physicochemical descriptors derived from RDKit, PaDEL, and Mordred, which summarize global molecular properties. The modality-specific embeddings are subsequently integrated through a fusion network that learns complementary structure– property relationships from these heterogeneous molecular representations.

Beyond predictive performance, an important objective of this work is to improve the interpretability and reliability of ADMET prediction models. To this end, DMPKformer incorporates modality-specific attention mechanisms for substructure-level interpretation and a latent-space out-of-distribution (OOD) confidence estimation framework to identify regions of reliable model operation. This combination of multimodal representation learning, interpretability, and reliability-aware prediction distinguishes DMPKformer from purely performance-driven ADMET benchmarking models.

## 2. Methodology

### 2.1 Data Sources

All datasets used in this study were obtained from the Therapeutics Data Commons (TDC), a standardized benchmark suite for machine learning in drug discovery. TDC provides a diverse collection of datasets spanning multiple ADMET (Absorption, Distribution, Metabolism, Excretion, and Toxicity) endpoints, along with predefined data splits to facilitate consistent and reproducible evaluation [13].

The use of these standardized splits ensures consistency with prior benchmarks and enables fair comparison across different modelling approaches and the distribution of the number of datapoints across various ADMET endpoints is graphically depicted as below in Fig. 1

**Fig. 1.**
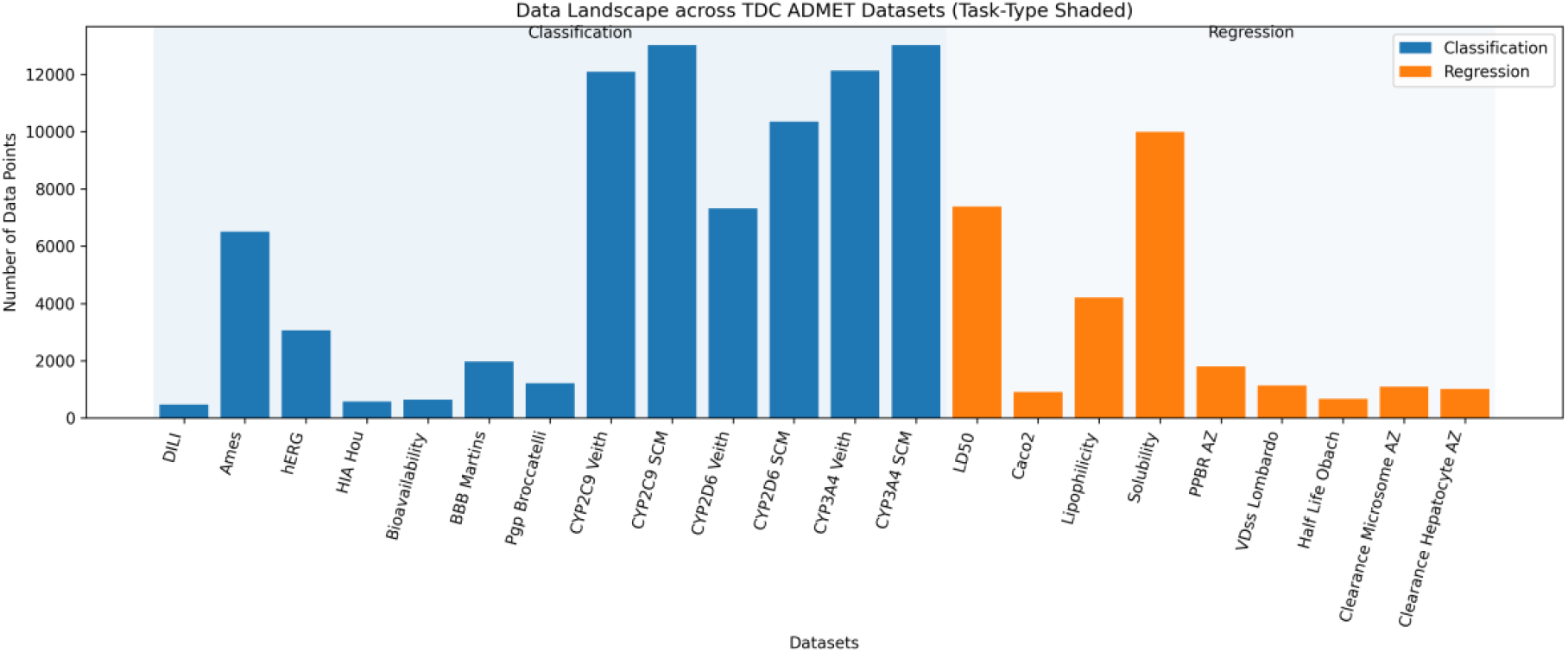
Data distribution of TDC across various ADMET endpoints

**Fig. 2.**
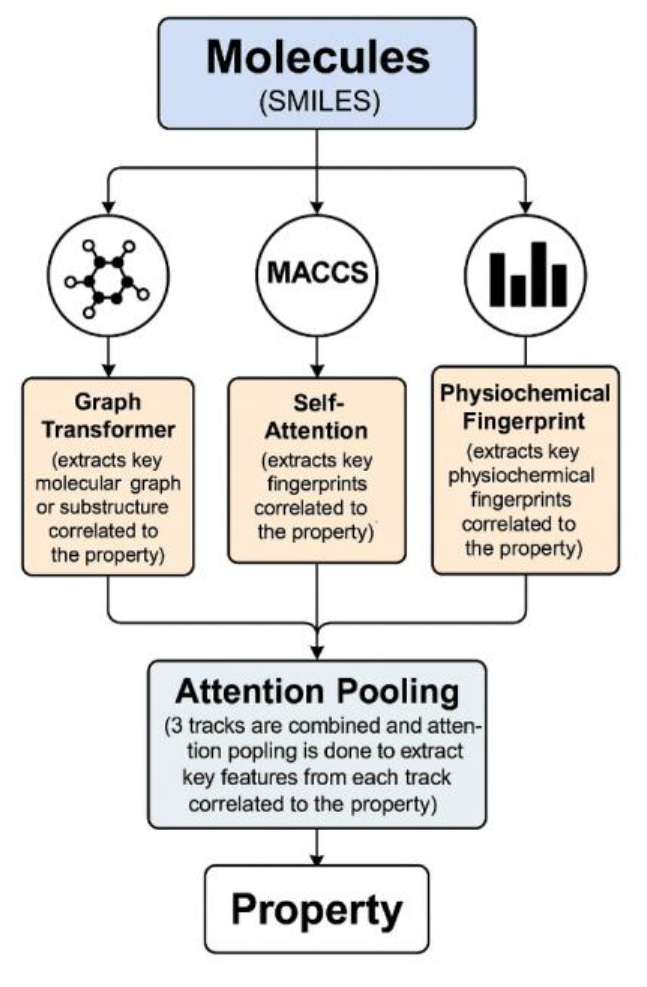
Overall architectural design of DMPKformer

### 2.2 Model Architecture

The proposed architecture consists of three independent modality-specific subnetworks and a final multimodal fusion network which is graphically depicted as below

#### 2.2.1 MACCS Fingerprint Track Architecture

The MACCS track is designed to extract expressive, task-adaptive representations from fixed-length binary molecular fingerprints, specifically the MACCS structural keys. Each molecule is represented as a 166-bit binary vector, where each bit encodes the presence or absence of a predefined chemical substructure or functional motif. While MACCS fingerprints are traditionally used with linear or shallow models, their fixed semantics and compact dimensionality make them well suited for contextual modelling via attention mechanisms. The architectural details of the MACCS self-attention transformer within the DMPKformer framework is given in Fig.3

**Fig. 3.**
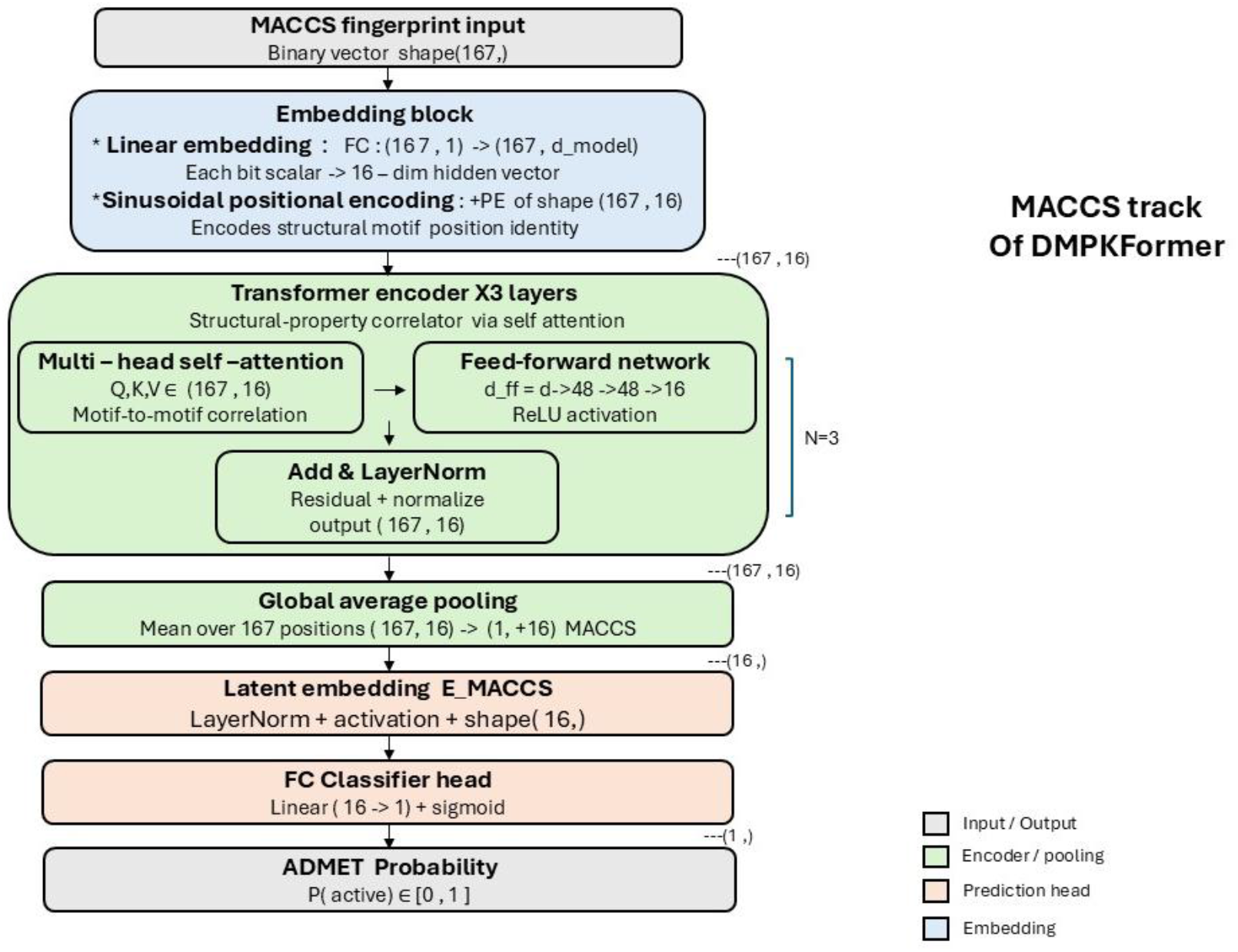
Architectural details of MACCS transformer

##### Input Representation and Embedding

The raw MACCS fingerprint, a binary vector of length 166, is first reshaped into a sequence of length 166 with a single feature channel per position. Each bit position is treated analogously to a token in a sequence, enabling the model to learn interactions between structural keys. A learnable linear embedding layer maps each scalar bit value into a continuous embedding space of dimension *d*_model_. This projection allows the network to represent both the presence of a structural feature and its contextual role relative to other features.

To preserve positional identity, since each MACCS bit corresponds to a specific chemical key, a fixed sinusoidal positional encoding is added to the embedded sequence. This ensures that the Transformer encoder can distinguish between different fingerprint indices while remaining invariant to batch ordering.

##### Transformer Encoder for Inter-Bit Dependency Modelling

The embedded fingerprint sequence is processed by a stack of Transformer encoder blocks, each consisting of a multi-head self-attention module followed by a position-wise feed-forward network. Unlike conventional fingerprint models that treat each bit independently or aggregate them linearly, self-attention enables the MACCS track to model higher-order dependencies between structural keys. This is particularly important in cheminformatics, where the predictive relevance of a substructure often depends on the presence or absence of other motifs elsewhere in the molecule.

Multi-head attention allows the model to attend to multiple interaction patterns in parallel, capturing diverse relationships such as co-occurrence of functional groups, mutually exclusive fragments, or long-range structural complementarities. The feed-forward sublayers further transform these context-enriched representations, while residual connections and normalization stabilize training and preserve information flow across layers.

##### Global Attention Pooling and Feature Attribution

Following the Transformer encoder, a global attention pooling mechanism aggregates the per-bit representations into a fixed-length molecular vector. This pooling layer learns a gating function that assigns an importance weight to each fingerprint position, effectively performing a soft selection over MACCS keys. The gated representations are then summed to produce a compact, molecule-level descriptor.

In addition to pooling, the attention weights are retained as per-bit importance scores, providing interpretability by highlighting which structural keys contribute most strongly to the model’s prediction. This aligns well with the interpretability expectations of fingerprint-based models in medicinal chemistry.

##### Latent Embedding and Prediction Head

The pooled molecular representation is passed through a normalization and non-linear activation stage to produce a latent embedding vector, denoted *E*_MACCS_. This embedding serves as a task-conditioned fingerprint representation that captures both local structural information and global context learned through attention. *E*_MACCS_can be used independently for downstream analysis, similarity comparisons, or as an input to multimodal fusion architectures.

For prediction, the latent embedding is processed by a dense feed-forward head composed of one or more fully connected layers with non-linear activation and dropout regularization. The final output layer produces a single logit, which is passed through a sigmoid activation to yield a probability for binary ADMET classification tasks. Training is performed using a weighted binary cross-entropy objective to account for class imbalance commonly observed in ADMET datasets.

#### 2.2.2 Molecular Graph Track Architecture

The Graph Track is designed to learn chemically grounded molecular representations directly from molecular structure, encoded as graphs derived from SMILES strings. Unlike fixed fingerprints, graph-based representations preserve explicit atomic connectivity and local chemical environments, allowing the model to reason about molecular structure in a manner that is closely aligned with physical chemistry. The architecture details of the Graph Attention Transformer adapted for molecular property prediction task within the DMPKformer framework is given in Fig. 4

**Fig. 4.**
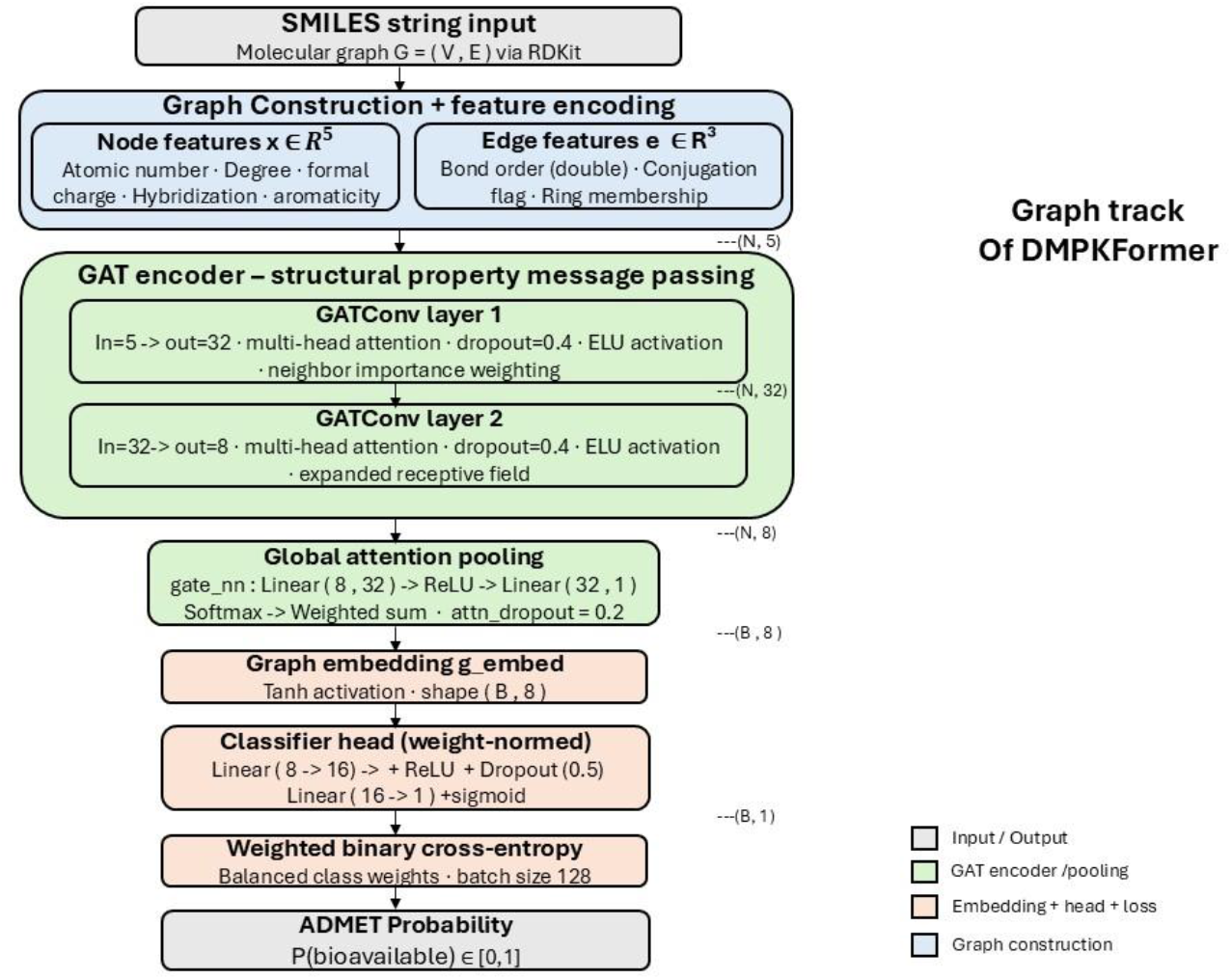
Architectural details of Graph Attention Transformer for molecular property prediction

##### Graph Construction and Feature Encoding

Each molecule is represented as a labelled graph *G* = (*V, E*), where nodes *V*correspond to atoms and edges *E*correspond to chemical bonds. Atom-level features are constructed using numerical descriptors that capture intrinsic chemical properties, such as atomic number, formal charge, degree, hybridization state, valence, aromaticity, and ring membership. These features are encoded as continuous or categorical values rather than one-hot vectors, enabling compact representations and smoother optimization.

Bond-level features encode the nature of chemical interactions between atoms, including bond order (single, double, triple), aromaticity, conjugation, and ring status. Together, these atom and bond descriptors provide a rich, physics-informed representation of molecular structure while remaining invariant to atom ordering and SMILES canonicalization.

##### Graph Attention Network for Local and Contextual Message Passing

The molecular graphs are processed using a Graph Attention Network (GAT), which performs message passing between neighbouring atoms while learning to weight the relative importance of different chemical interactions. In each GAT layer, node embeddings are updated by aggregating features from neighbouring nodes, where the contribution of each neighbour is modulated by a learned attention coefficient. These coefficients are computed as a function of both the source and target node representations (and, optionally, edge features), allowing the network to prioritize chemically relevant interactions such as polar bonds, aromatic systems, or functional group connectivity.

Multi-head attention is employed to enable the model to capture diverse interaction patterns simultaneously. Each attention head can focus on different aspects of the local chemical environment, such as steric effects, electronic properties, or substructure motifs. The outputs of multiple heads are concatenated or averaged to form the updated node embeddings, improving representational capacity and robustness.

By stacking multiple GAT layers, the model progressively expands the receptive field of each atom, allowing information to propagate across larger molecular substructures. This hierarchical message passing enables the Graph Track to capture both local atomic environments and higher-order structural motifs such as rings, scaffolds, and functional group arrangements.

##### Global Attention Pooling and Molecular Representation

After graph-level message passing, node embeddings are aggregated into a fixed-length molecular representation using a global attention pooling mechanism. Rather than simply averaging or summing node features, attention pooling learns a weighting over atoms that reflects their relative importance to the prediction task. This allows the model to emphasize chemically salient atoms—such as reactive centres, pharmacophores, or key substituents— while down-weighting less informative regions of the molecule.

The pooling operation produces a single vector representation for each molecule that integrates information across the entire graph while retaining interpretability via learned attention weights. These weights can be inspected post hoc to identify which atoms or substructures most strongly influenced the model’s prediction, supporting mechanistic interpretation and hypothesis generation.

##### Prediction Head and Latent Graph Embedding

The pooled molecular graph representation is passed through a dense feed-forward prediction head composed of fully connected layers with non-linear activations and regularization. The final layer outputs a single logit, which is transformed via a sigmoid activation to produce a probability for binary classification tasks such as ADMET property prediction.

The intermediate pooled graph vector, extracted prior to the final prediction layer, is denoted as *E*_Graph_. This latent embedding captures a task-conditioned, structure-aware representation of the molecule that reflects both atomic-level chemistry and global molecular context. *E*_Graph_can be used independently for downstream analyses, similarity comparisons, or integration with other molecular modalities.

#### 2.2.3 Physicochemical Descriptor Track

Physicochemical and topological descriptors are computed from RDKit, PaDEL, and Mordred, providing high-dimensional continuous features summarizing global molecular properties. These features are fed into a tanh-activated multilayer perceptron (MLP) network that captures nonlinear dependencies among descriptors. An embedding vector from an internal dense layer serves as E_Desc, representing the learned descriptor-level representation. The architectural details of the descriptor track is provided in Fig. 5

**Fig. 5.**
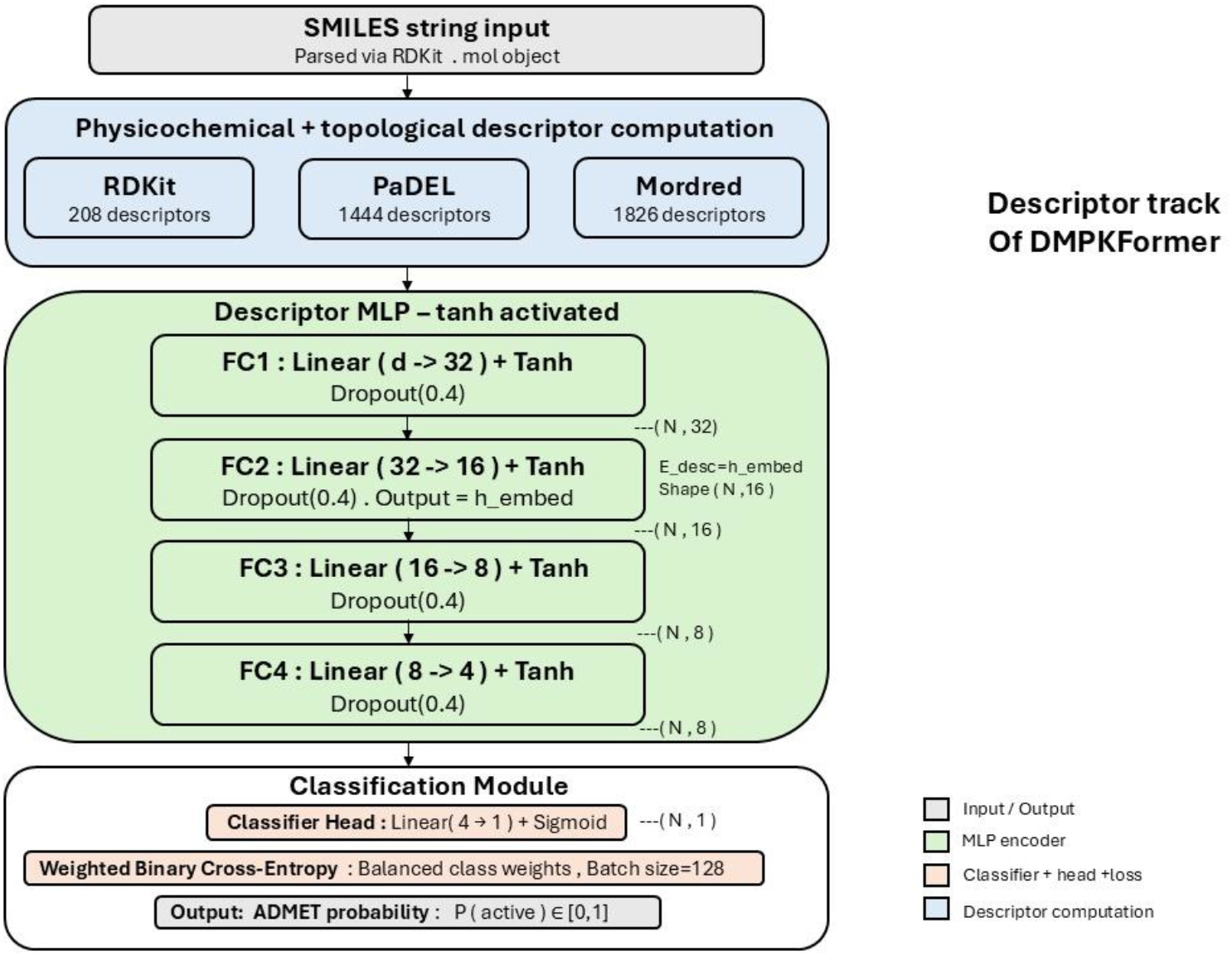
Architectural details of physicochemical descriptor track in the DMPKformer framework

### 2.3 Phase-wise Training Strategy

Each subnetwork is independently trained for binary classification using a dataset of molecules annotated with the target ADMET label. Binary cross-entropy loss is used for optimization, and L2 regularization and dropout are applied to prevent overfitting. Once individual models converge, intermediate embeddings E_MACCS, E_Graph, and E_Desc are extracted for all molecules.

These embeddings form the input to the fusion network, where they are concatenated and passed through a tanh-activated dense network that models inter-modality interactions. The final classification layer produces the unified ADMET prediction output. During this phase, the modality encoders remain frozen to preserve previously learned representations, and only the fusion network is trained.

In the section that follows we present and discuss the results obtained for the independent tracks and the combined track and compare them against state of art benchmarks and demonstrate improvement over state of art.

## 3 Results and Discussion

The following section presents the predictive performance of DMPKformer across multiple TDC benchmarks together with an analysis of the interpretability provided by the framework. In addition, we introduce a latent-space out-of-distribution (OOD) metric for TDC tasks that enables reliability-aware confidence estimation and identification of high-confidence regions of model operation.

### 3.1 Track-wise results of DMPKformer

The results from the MACCS self-attention transformer, Graph Attention transformer and the physicochemical descriptor track are presented below for the classifications and regression tasks in TDC in table 1a, 1b, 2a, 2b & 3a, 3b respectively.

**Table 1a.** MACCS track of DMPKformer on TDC classification tasks.

| Task | Metrics (↑ ) | Maccs Track |
| --- | --- | --- |
| DILI | AUROC | 0.912 |
| Ames | AUROC | 0.790 |
| HIA Hou | AUROC | 0.974 |
| Bioavailability | AUROC | 0.610 |
| hERG | AUROC | 0.830 |
| CYP2D6<br>SCM | AUPRC | 0.549 |
| CYP3A4<br>SCM | AUROC | 0.541 |
| CYP2C9<br>SCM | AUPRC | 0.448 |
| CYP2C9<br>Veith | AUPRC | 0.677 |
| CYP3A4<br>Veith | AUPRC | 0.791 |
| CYP2D6<br>Veith | AUPRC | 0.611 |
| BBB Martins | AUROC | 0.878 |
| Pgp<br>Broccatelli | AUROC | 0.890 |

**Table 1b.** MACCS track of DMPKformer on TDC regression tasks.

| Task | Metrics | Maccs Track |
| --- | --- | --- |
| Caco2 | MAE ↓ | 0.341 |
| LD50 | MAE ↓ | 0.629 |
| Lipophilicity | MAE ↓ | 0.710 |
| PPBR AZ | MAE ↓ | 13.430 |
| Solubility<br>AqSolDB | MAE ↓ | 1.700 |
| VDss Lombardo | Spearman Corr. ↑ | 0.530 |
| Half Life Obach | Spearman Corr. ↑ | 0.420 |
| Clearance<br>Microsome AZ | Spearman Corr. ↑ | 0.450 |
| Clearance<br>Hepatocyte AZ | Spearman Corr. ↑ | 0.300 |

**Table 2a.** Graph track of DMPKformer on TDC classification tasks.

| Task | Metrics (↑ ) | Graph Track |
| --- | --- | --- |
| DILI | AUROC | 0.698 |
| Ames | AUROC | 0.787 |
| HIA Hou | AUROC | 0.809 |
| Bioavailability | AUROC | 0.750 |
| hERG | AUROC | 0.758 |
| CYP2D6 SCM | AUPRC | 0.761 |
| CYP3A4 SCM | AUROC | 0.578 |
| CYP2C9 SCM | AUPRC | 0.332 |
| CYP2C9 Veith | AUPRC | 0.661 |
| CYP3A4 Veith | AUPRC | 0.754 |
| CYP2D6 Veith | AUPRC | 0.483 |
| BBB Martins | AUROC | 0.878 |
| Pgp Broccatelli | AUROC | 0.901 |

**Table 2b.** Graph track of DMPKformer on TDC regression tasks.

| Task | Metrics | Graph track |
| --- | --- | --- |
| Caco2 | MAE ↓ | 0.459 |
| LD50 | MAE ↓ | 0.731 |
| Lipophilicity | MAE ↓ | 0.737 |
| PPBR AZ | MAE ↓ | 9.244 |
| Solubility AqSolDB | MAE ↓ | 1.148 |
| VDss Lombardo | Spearman Corr. ↑ | 0.626 |
| Half Life Obach | Spearman Corr. ↑ | 0.620 |
| Clearance Microsome AZ | Spearman Corr. ↑ | 0.617 |
| Clearance Hepatocyte AZ | Spearman Corr. ↑ | 0.321 |

**Table 3a.** Descriptor track of DMPKformer on TDC classification tasks.

| Task | Metrics (↑ ) | Rdkit Track | Mordred track |
| --- | --- | --- | --- |
| DILI | AUROC | 0.881 | 0.909 |
| Ames | AUROC | 0.824 | 0.833 |
| HIA Hou | AUROC | 0.978 | 0.973 |
| Bioavailability | AUROC | 0.740 | 0.700 |
| hERG | AUROC | 0.865 | 0.862 |
| CYP2D6 SCM | AUPRC | 0.725 | 0.679 |
| CYP3A4 SCM | AUROC | 0.685 | 0.541 |
| CYP2C9 SCM | AUPRC | 0.462 | 0.385 |
| CYP2C9 Veith | AUPRC | 0.719 | 0.726 |
| CYP3A4 Veith | AUPRC | 0.827 | 0.830 |
| CYP2D6 Veith | AUPRC | 0.623 | 0.630 |
| BBB Martins | AUROC | 0.914 | 0.916 |
| Pgp Broccatelli | AUROC | 0.911 | 0.922 |

**Table 3b.**
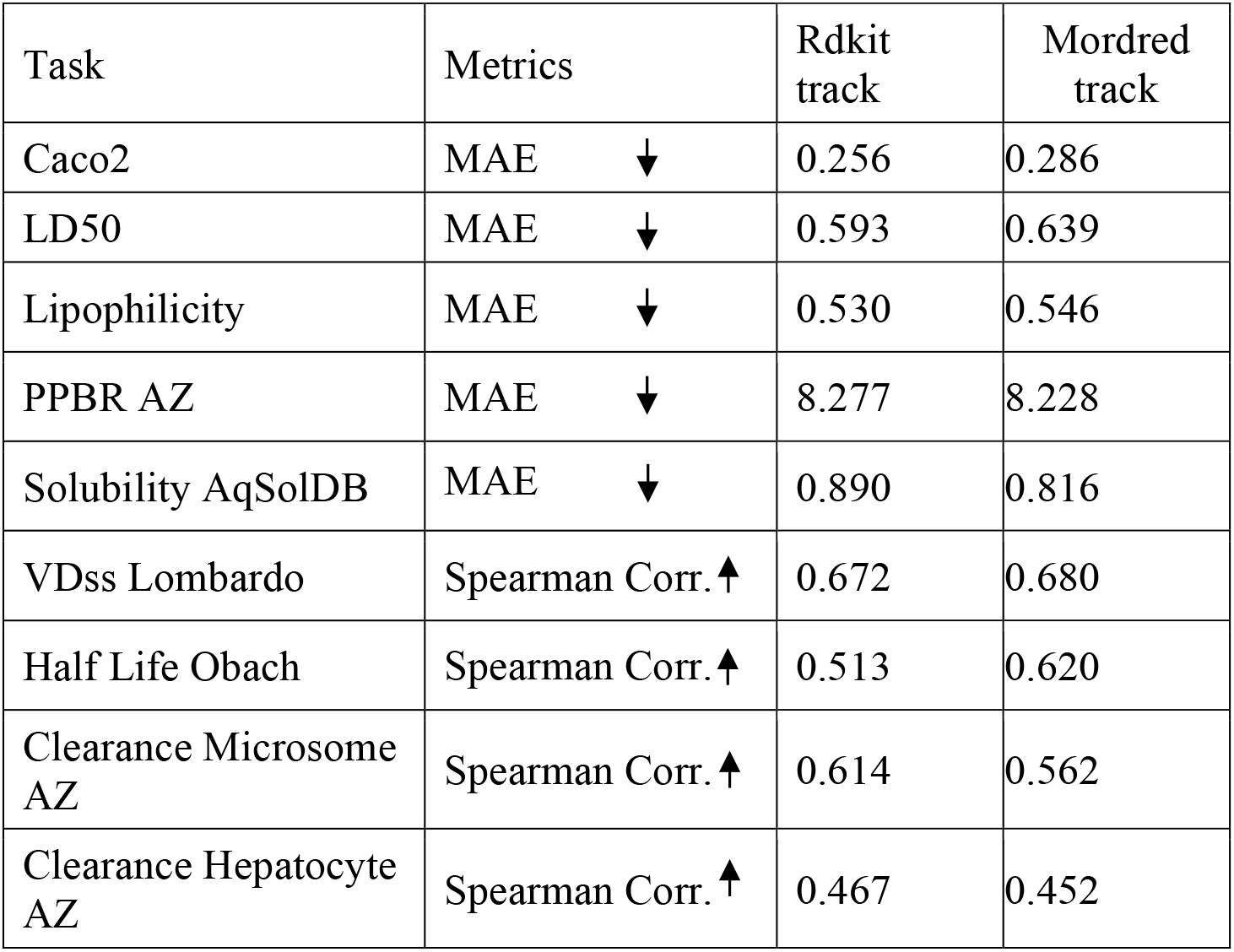
Descriptor track of DMPKformer on TDC regression tasks.

The central premise underlying DMPKformer is that each modality-specific branch captures a distinct and complementary view of molecular structure and behavior. The MACCS fingerprint branch encodes fragment-level structural motifs and predefined chemical patterns, the graph-based branch captures topology-aware atomic interactions and local chemical environments, while the physicochemical descriptor summarizes global molecular properties relevant to pharmacokinetic and toxicity behavior. Individually, each modality provides only a partial representation of molecular characteristics governing ADMET outcomes. However, the latent embeddings learned by the independent subnetworks encode complementary structural and physicochemical information that can be integrated to form a richer and more generalizable molecular representation.

By combining these modality-specific embeddings through a unified fusion network, DMPKformer learns higher-order cross-modal structure–property relationships that are not fully accessible through any individual representation alone. This multimodal integration improves the model’s ability to capture the complex and heterogeneous factors governing ADMET behavior across diverse molecular scaffolds. The performance of the combined multimodal DMPKformer architecture on TDC classification and regression benchmarks is presented in Tables 4a and 4b, respectively.

**Table 4a.**
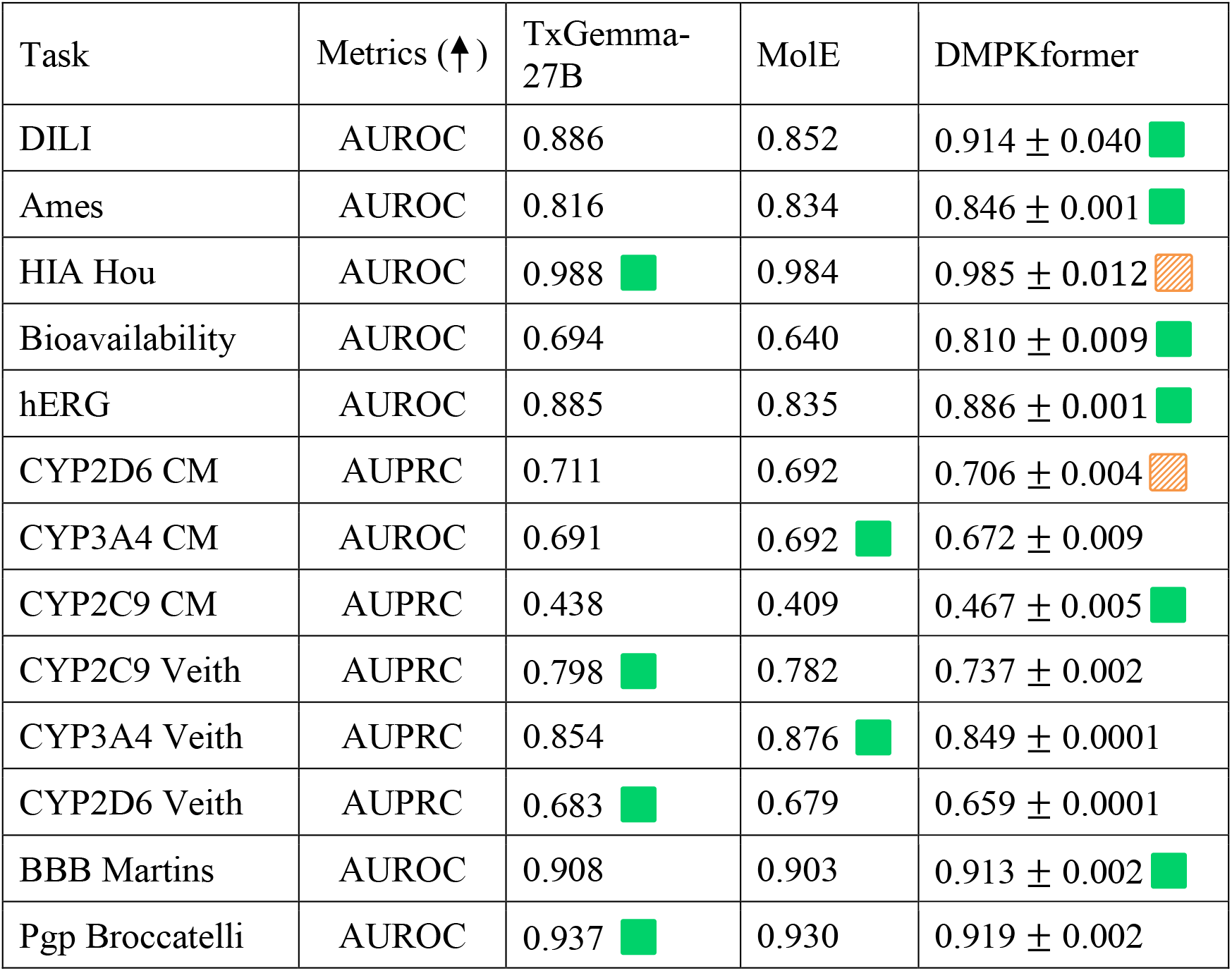
DMPKformer on TDC classification tasks.

| Task | Metrics (↑) | TxGemma-27B | MolE | DMPKformer |
| --- | --- | --- | --- | --- |
| DILI | AUROC | 0.886 | 0.852 | $0.914 \pm 0.040$ |
| Ames | AUROC | 0.816 | 0.834 | $0.846 \pm 0.001$ |
| HIA Hou | AUROC | 0.988 | 0.984 | $0.985 \pm 0.012$ |
| Bioavailability | AUROC | 0.694 | 0.640 | $0.810 \pm 0.009$ |
| hERG | AUROC | 0.885 | 0.835 | $0.886 \pm 0.001$ |
| CYP2D6 CM | AUPRC | 0.711 | 0.692 | $0.706 \pm 0.004$ |
| CYP3A4 CM | AUROC | 0.691 | 0.692 | $0.672 \pm 0.009$ |
| CYP2C9 CM | AUPRC | 0.438 | 0.409 | $0.467 \pm 0.005$ |
| CYP2C9 Veith | AUPRC | 0.798 | 0.782 | $0.737 \pm 0.002$ |
| CYP3A4 Veith | AUPRC | 0.854 | 0.876 | $0.849 \pm 0.0001$ |
| CYP2D6 Veith | AUPRC | 0.683 | 0.679 | $0.659 \pm 0.0001$ |
| BBB Martins | AUROC | 0.908 | 0.903 | $0.913 \pm 0.002$ |
| Pgp Broccatelli | AUROC | 0.937 | 0.930 | $0.919 \pm 0.002$ |

**Table 4b.**
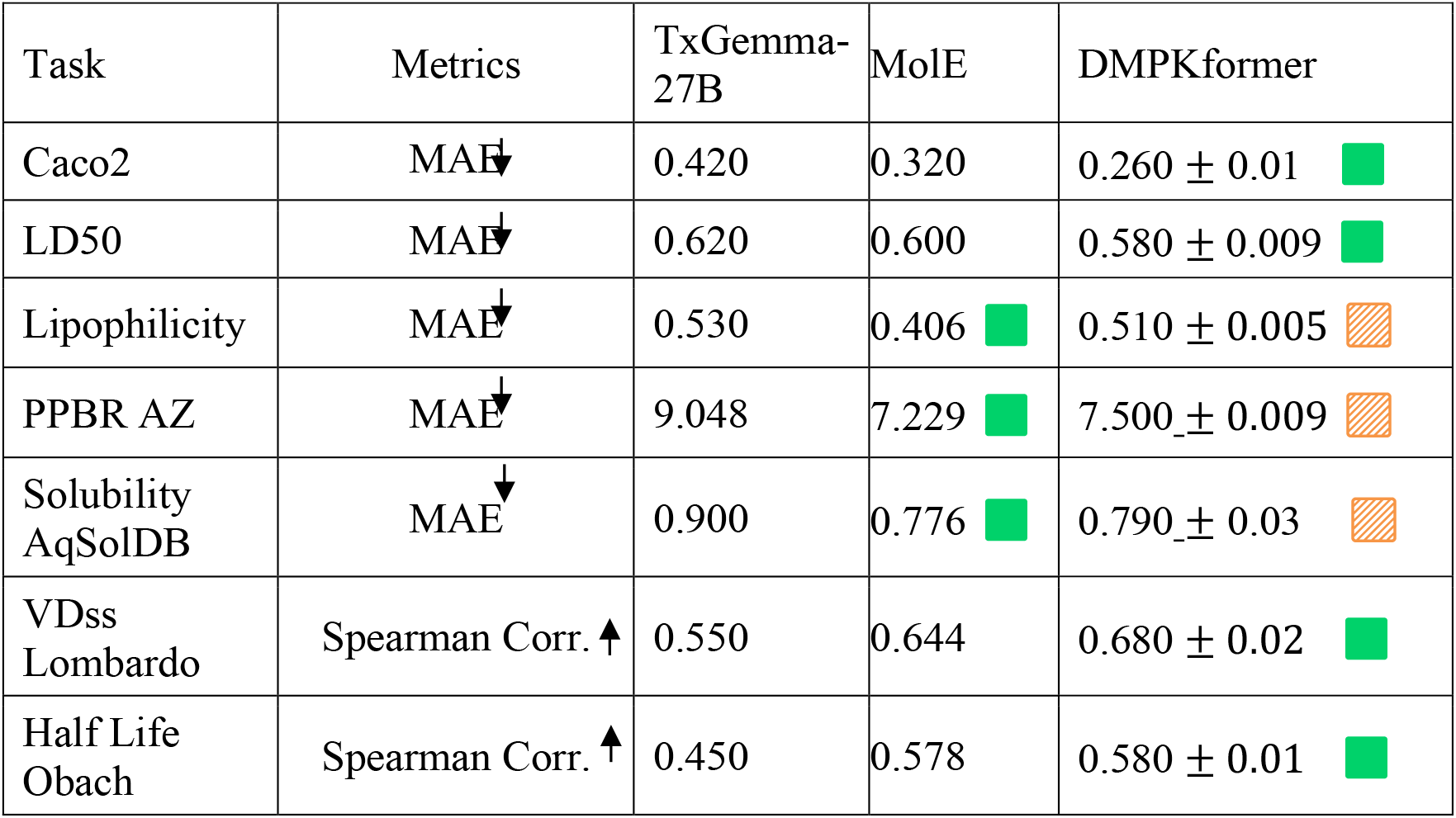
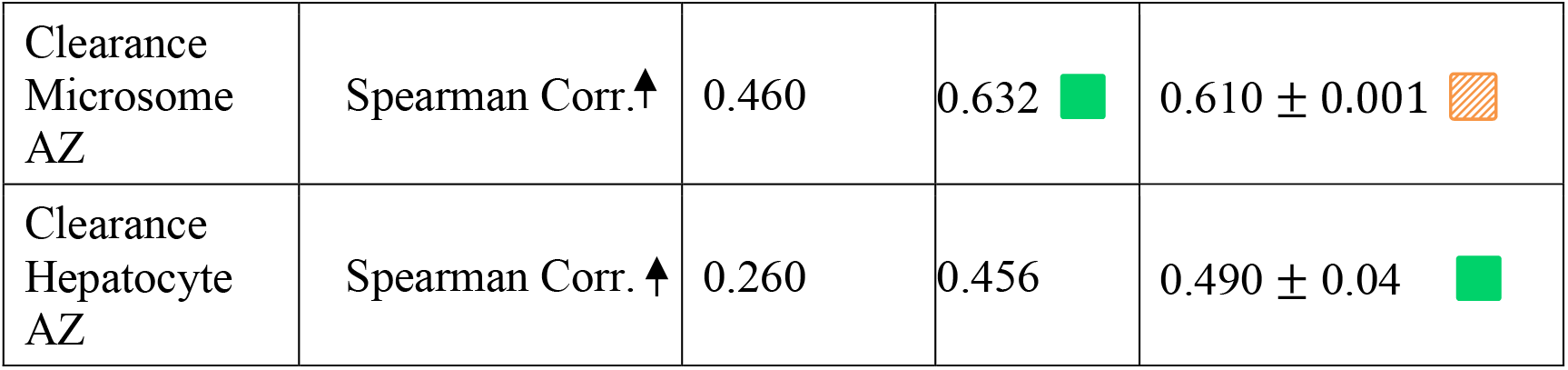
DMPKformer on TDC regression tasks.

| Task | Metrics | TxGemma-27B | MolE | DMPKformer |
| --- | --- | --- | --- | --- |
| Caco2 | MAE ↓ | 0.420 | 0.320 | $0.260 \pm 0.01$ |
| LD50 | MAE ↓ | 0.620 | 0.600 | $0.580 \pm 0.009$ |
| Lipophilicity | MAE ↓ | 0.530 | 0.406 | $0.510 \pm 0.005$ |
| PPBR AZ | MAE ↓ | 9.048 | 7.229 | $7.500 \pm 0.009$ |
| Solubility AqSolDB | MAE ↓ | 0.900 | 0.776 | $0.790 \pm 0.03$ |
| VDss Lombardo | Spearman Corr. ↑ | 0.550 | 0.644 | $0.680 \pm 0.02$ |
| Half Life Obach | Spearman Corr. ↑ | 0.450 | 0.578 | $0.580 \pm 0.01$ |
| Clearance<br>Microsome<br>AZ | Spearman Corr.↑ | 0.460 | 0.632 ■ | $0.610 \pm 0.001$ ▨ |
| Clearance<br>Hepatocyte<br>AZ | Spearman Corr. ↑ | 0.260 | 0.456 | $0.490 \pm 0.04$ ■ |

The performance of DMPKformer was compared against recent literature benchmark models, including TxGemma [14] and MolE [15], developed by groups such as DeepMind and Recursion, respectively. Across multiple TDC classification and regression tasks, DMPKformer achieved competitive or superior performance on several important ADMET endpoints, including DILI, Ames mutagenicity, hERG inhibition, and bioavailability prediction. Performance improvements over competing methods are highlighted in green in Tables 4a and 4b.

To further analyze overall model behavior across tasks, the comparative rankings of DMPKformer, MolE, and TxGemma are summarized in Fig. 6a and Fig. 6b. Fig. 6a illustrates the number of tasks on which each model achieves the highest performance, demonstrating the strong overall competitiveness of DMPKformer across diverse ADMET endpoints. Fig. 6b presents the distribution of model ranks across tasks, where DMPKformer exhibits a lower median rank and a narrower spread toward top-performing positions, indicating more consistent performance across heterogeneous datasets. In contrast, MolE demonstrates moderate rank consistency, while TxGemma exhibits greater variability across tasks.

**Fig 6a.**
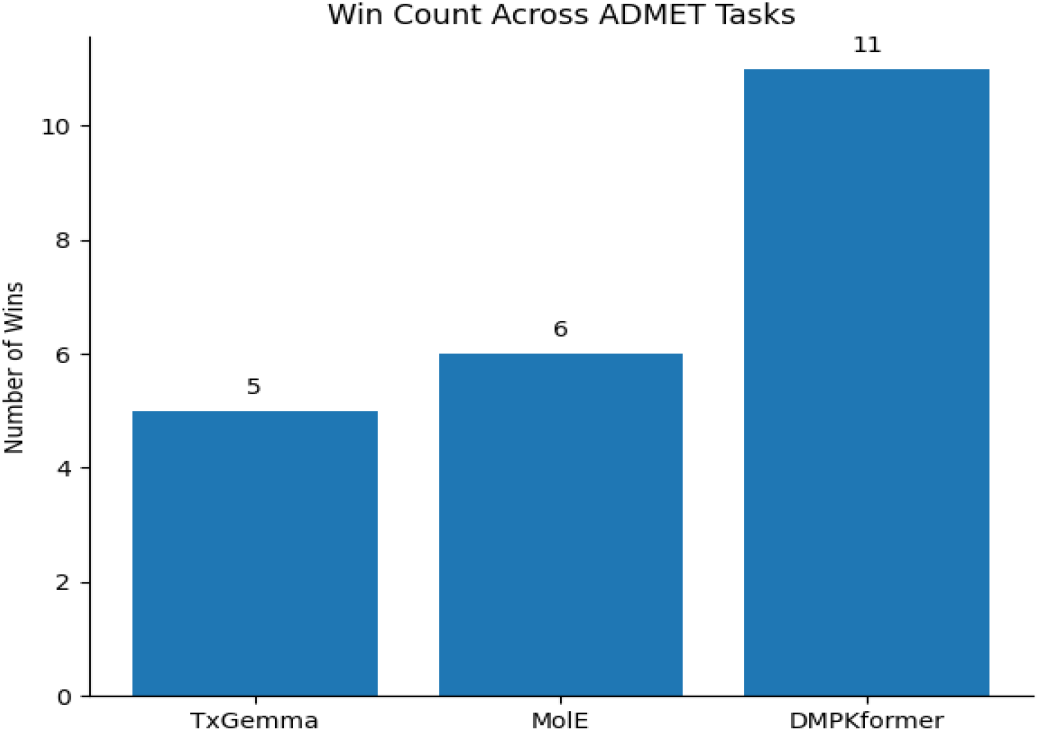
DMPKformer state of art demonstration

**Fig 6b.**
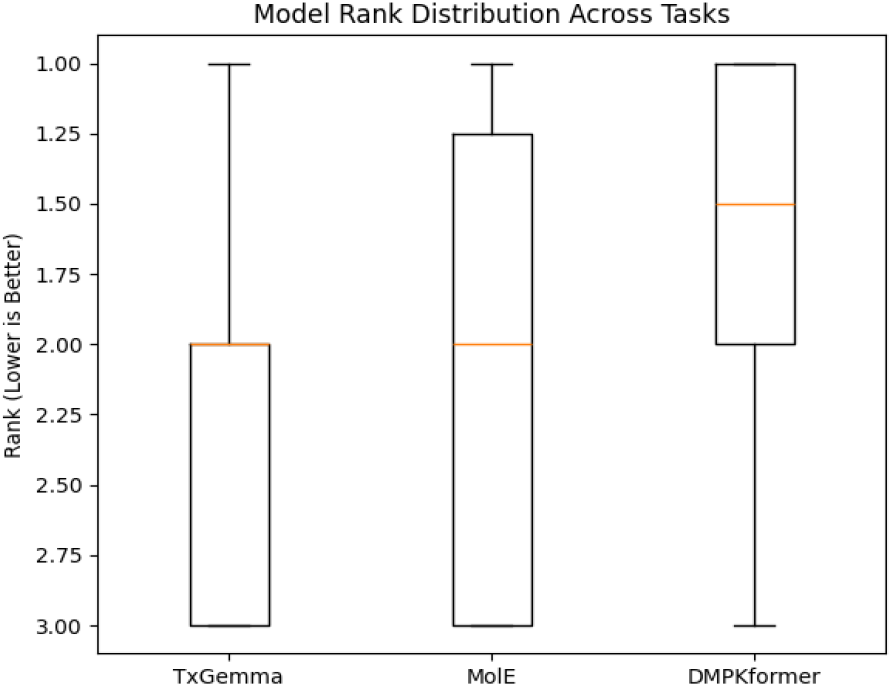
DMPKformer state of art demonstration

### 3.4 Interpretability and Modularity

The modular training framework enables direct visualization and interpretation of the latent embedding spaces learned by each modality-specific branch. Attention mechanisms within the MACCS and graph-based branches provide substructure-level interpretability by highlighting molecular fragments and atomic neighborhoods that contribute strongly to ADMET predictions. In parallel, descriptor-based sensitivity analysis elucidates the influence of global physicochemical properties on classification outcomes.

For example, structural motifs associated with Ames mutagenicity, including polycyclic aromatic hydrocarbons such as phenanthrene and anthracene, were assigned high node-importance scores within the DMPKformer interpretability framework, consistent with previously reported mutagenicity-associated structural alerts [16,17], as illustrated in Figure 7.

**Fig 7.**
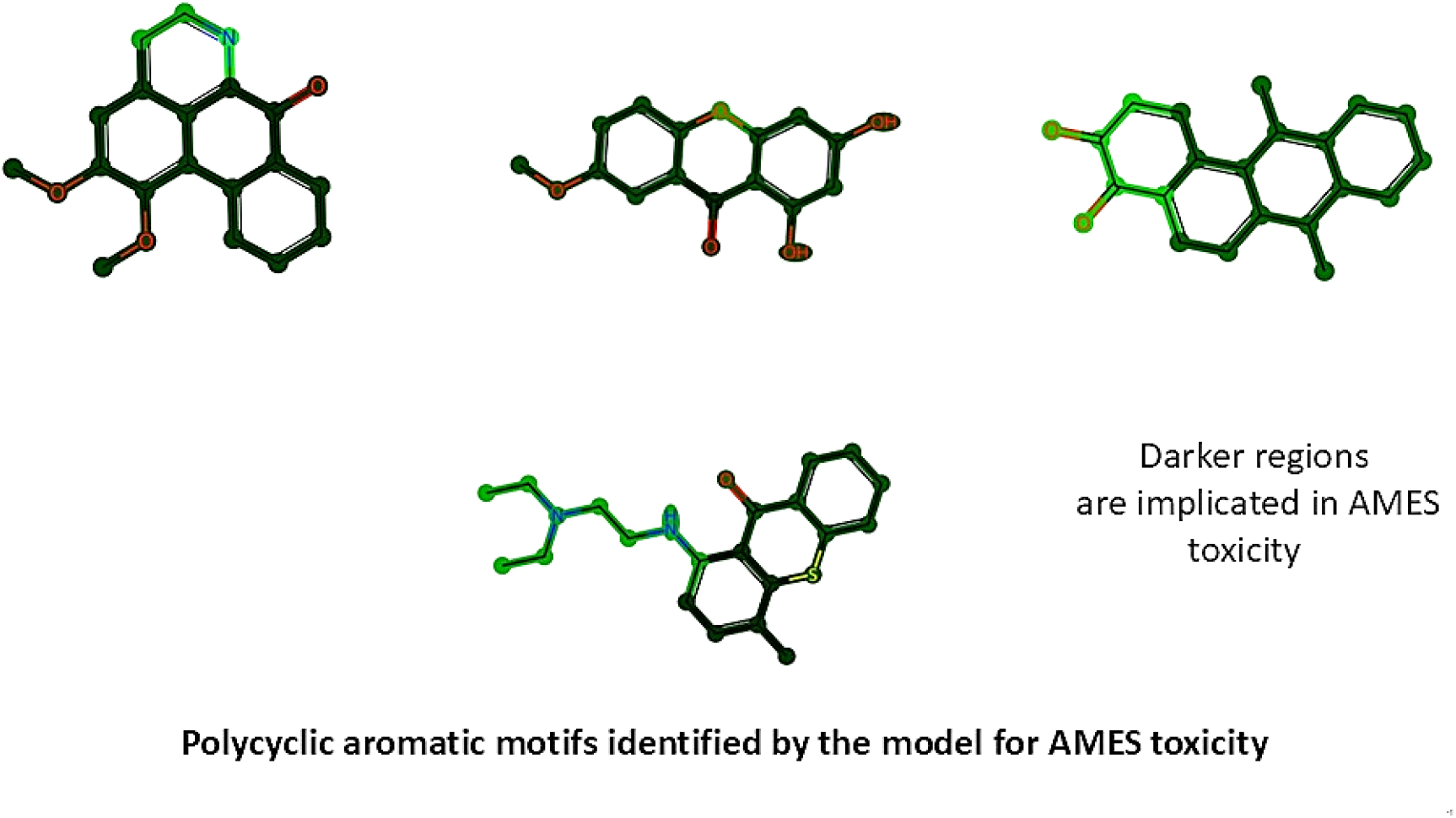
Interpretability in DMPKformer

## 4. Latent space derived Out-of-Distribution vs test accuracy to define high confidence region of operation of DMPKformer

To define a reliability-aware operating region for DMPKformer, we introduce a latent-space derived out-of-distribution (OOD) metric [18–20] based on PCA-derived similarity analysis of the learned molecular embeddings. For a given test molecule, the OOD score quantifies the deviation of its latent representation from the distribution of training embeddings. The relationship between prediction accuracy and OOD score is analysed by plotting test-set accuracy against the corresponding OOD values.

As shown in Figure 8 for the Ames mutagenicity dataset, prediction accuracy decreases progressively with increasing OOD score, indicating that model reliability deteriorates as test molecules become more dissimilar to the training distribution. A high-confidence operating region is observed within an OOD range of approximately 0–0.12, where prediction accuracy remains above 80%, while accuracy declines below 65% for OOD values greater than 0.2. These results demonstrate that the proposed latent-space OOD metric provides a meaningful indicator of prediction reliability and operational confidence for ADMET prediction tasks.

**Fig. 8.**
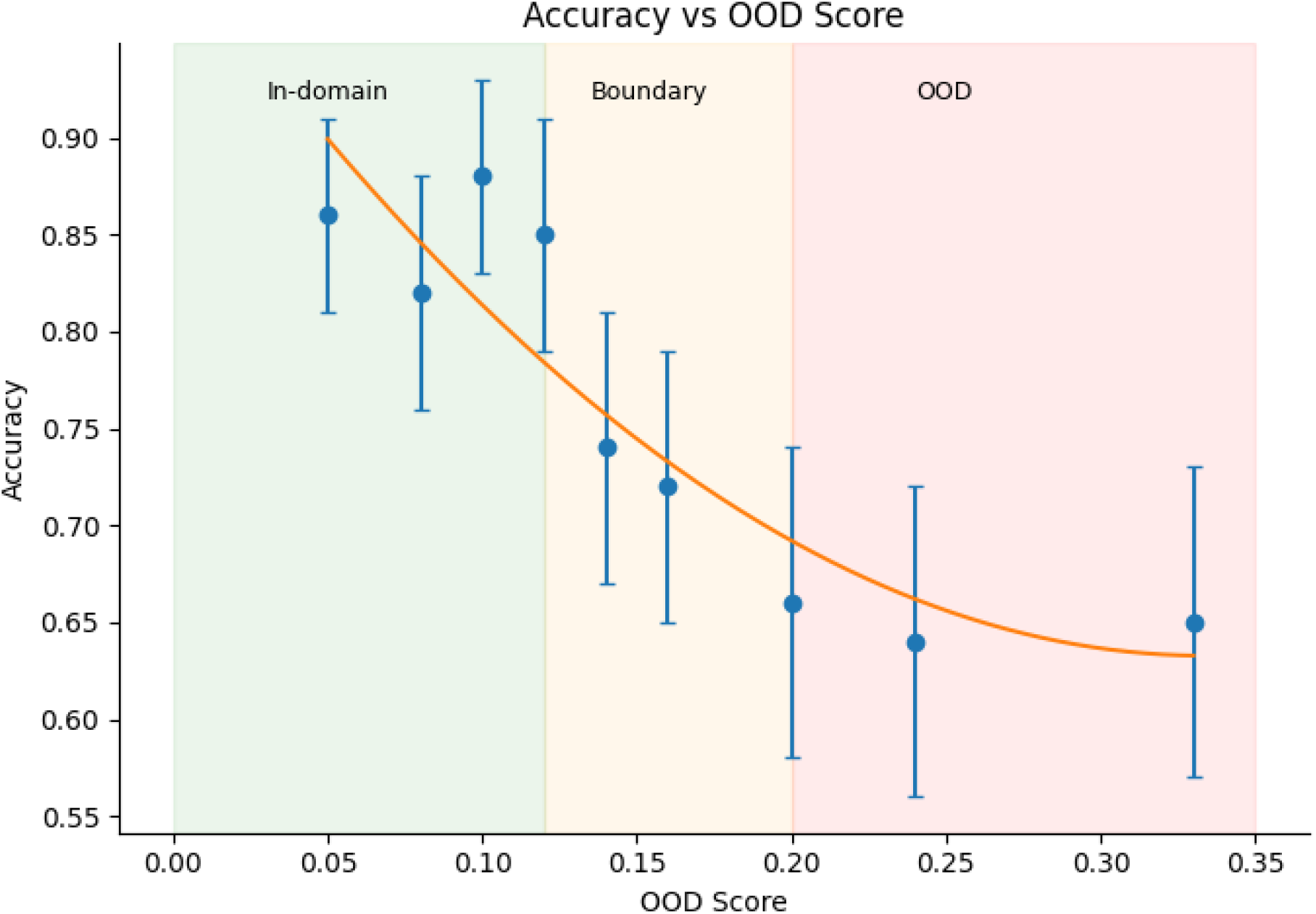
Out-Of-Distribution (OOD) score vs Test accuracy

The OOD-based confidence analysis was further extended across the complete TDC benchmark suite to characterize the operational confidence regions of DMPKformer for both classification and regression tasks. The resulting in-domain and out-of-domain performance statistics are summarized in Tables 5a and 5b.

**Table 5a.**
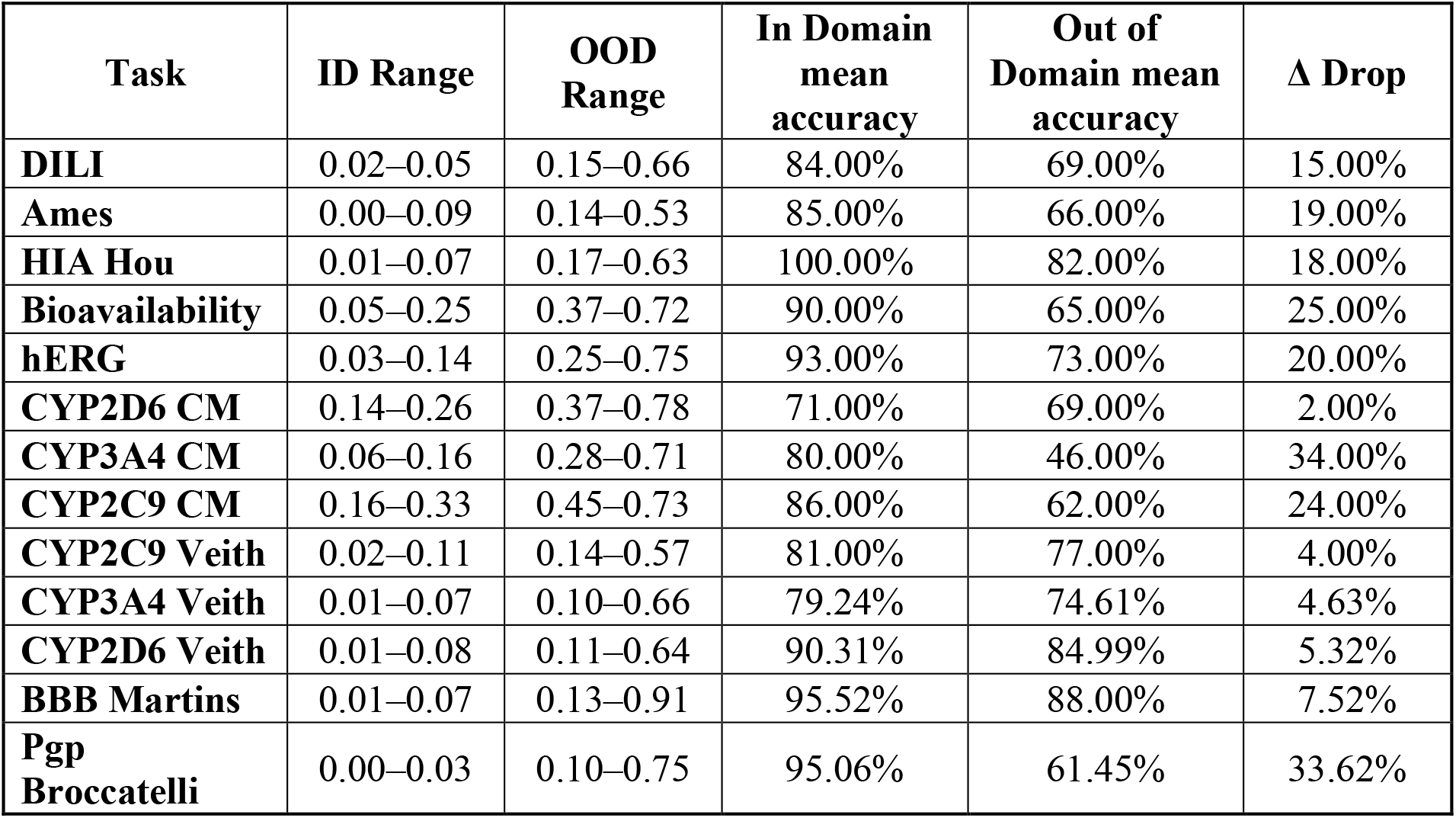
DMPKFormer’s operating region of confidence on TDC classification dataset defined based on OOD metric.

**Table 5b.** DMPKFormer’s operating region of confidence on TDC regression dataset defined based on OOD metric.

| Task | ID Range | OOD Range | In Domain MAE | Out of Domain MAE | $\Delta$ Drop |
| --- | --- | --- | --- | --- | --- |
| **Caco2** | 0.07–0.23 | 0.32–0.68 | 0.27 | 0.29 | 0.02 |
| **LD50** | 0.07–0.20 | 0.26–0.74 | 0.49 | 0.73 | 0.24 |
| **Lipophilicity** | 0.09–0.21 | 0.28–0.75 | 0.48 | 0.54 | 0.06 |
| **PPBR AZ** | 0.05–0.12 | 0.17–0.64 | 5.78 | 9.59 | 3.81 |
| **Solubility AqSolDB** | 0.02–0.74 | -- | 0.89 | No significant out of domain observed | -- |
| **VDss Lombardo** | 0.07–0.17 | 0.22–0.78 | 0.76 | 0.67 | 0.09 |
| **Half Life Obach** | 0.02–0.11 | 0.15–1.00 | 0.65 | 0.50 | 0.15 |
| **Clearance Microsome AZ** | 0.10–0.69 | -- | 0.58 | No significant out of domain observed | -- |
| **Clearance Hepatocyte AZ** | 0.08–0.81 | -- | 0.38 | No significant out of domain observed | -- |

## 5. Conclusion and Future Scope

In this study, we presented DMPKformer, a multimodal deep learning framework for ADMET property prediction that integrates MACCS fingerprints, graph-based molecular representations, and physicochemical descriptors through specialized subnetworks and a unified fusion strategy. Each modality-specific encoder captures complementary levels of chemical abstraction, including fragment-level structural patterns, topology-aware atomic interactions, and global physicochemical properties, enabling the model to learn richer and more generalizable molecular representations.

The predictive framework of DMPKformer is designed as a modular and interpretable framework that combines complementary molecular representations with reliability-aware confidence estimation through latent-space out-of-distribution (OOD) analysis. This integration of predictive performance, interpretability, and operational confidence estimation enhances the practical applicability of the framework for real-world drug discovery workflows as a valuable tool for early-stage virtual screening, toxicity assessment, and lead optimization in computational drug discovery pipelines.

Future work will focus on incorporating additional feature tracks, advanced cross-modal fusion mechanisms which improves crosstalk across feature tracks improving learning, and uncertainty-awareness learning strategies to further improve generalization, robustness, and applicability across broader molecular property prediction tasks.

